# Pollen-feeding delays reproductive senescence and maintains toxicity of *Heliconius erato*

**DOI:** 10.1101/2023.01.13.523799

**Authors:** Erika C. Pinheiro de Castro, Josie McPherson, Glennis Jullian, Anniina L. K. Mattila, Søren Bak, Stephen H. Montgomery, Chris Jiggins

## Abstract

Dietary shifts may act to ease energetic constraints and allow organisms to optimise life-history traits. *Heliconius* butterflies differ from other nectar-feeders due to their unique ability to digest pollen, which provides a reliable source of amino acids to adults. Pollen-feeding has been associated with prolonged adult lifespan and increased fertility, yet there is a lack of empirical data demonstrating how pollen consumption influences key fitness traits, including chemical defences and adult body weight, as well as fertility over their elongated lifespan. Here, we investigated the effect of pollen-feeding on fertility, weight and chemical defences, as well as offspring defences, controlling for butterfly age and sex. Recently emerged *Heliconius erato* butterflies of similar size were fed for 14 or 45 days on one of three diets: sugar solution only, or sugar solution replenished with either amino acid supplement or pollen. At the end of the experiment, oviposition assays were performed to evaluate fertility, and afterwards all butterflies and eggs were weighed and used for quantification of cyanogenic glucosides (CG). We found that there is an age-specific and sex-specific effect of pollen-feeding on butterfly weight, with both the sugar-only and amino-acid supplement diets reducing the weight of old females (45d), but not young females (14d) or males of any age. Females fed only sugar significantly reduced their egg-laying through adulthood, whereas females that had access to pollen maintained their fertility. Diet had a significant effect on the maintenance of the chemical defence of females, but not males. Curiously, even though females that have access to pollen were heavier, more toxic and laid more eggs, this did not translate into improvements in offspring defences, as eggs from butterflies of all ages and diet treatments had similar CG content. Our results emphasise the importance of controlling for age-specific and sex-specific effects in studies of life-history evolution and demonstrate that dietary novelty can relax energetic constraints.

## Introduction

Survival and reproductive success are the two major components of Darwinian fitness, and as with all life-history traits, they are under strong selective pressures. Nevertheless, species cannot evolve to live forever and reproduce continuously (there are no “Darwinian demons” (Law, 1979)) due to physiological and energetic constraints that create trade-offs between life-history traits (Healy, Ezard, Jones, Salguero-Gómez, & Buckley, 2019). Yet, dietary shifts may ease energetic constraints, which could in turn allow organisms to optimise multiple fitness traits simultaneously (Swanson et al., 2016).

One striking case of dietary innovation is provided by the pollen feeding *Heliconius* butterflies (Gilbert 1972; Young and Montgomery 2020). Butterflies typically require water and sugars during adulthood, which can be acquired either from rotten fruits (fruit-feeders) or nectar produced by flowers (nectar-feeders) (Krenn, 2008). Butterflies of the *Heliconius* genus differ from other nectar-feeders due to their ability to additionally collect and digest pollen while feeding on nectar (Gilbert 1972; Young and Montgomery 2020). Although many insects can eat pollen (*e*.*g*. bees as well as some beetles, sawflies, mirids, thrips, flies and moths) (Wäckers, Romeis, & Van Rijn, 2007), *Heliconius* are the only butterflies known to actively collect and digest pollen grains. This is probably explained by the necessity of specific adaptations for mechanical and chemical digestion of pollen to make its nutrients available for absorption (Johnson & Nicolson, 2001).

A number of adaptations were probably necessary to allow *Heliconius* to digest pollen (Harpel et al., 2015; Smith et al., 2016; Smith et al., 2018; Cicconardi et al., 2023). Pollen grains collected in the elongated proboscis of these butterflies are humidified with salivary secretions, aided by the co-option of a “grooming behaviour” (coiling and uncoiling of the proboscis for some minutes to hours) (Gilbert 1972; Krenn et al. 2009; Hikl and Krenn 2011). Pollen-feeding is not observed in other genera of the Heliconiini tribe and arose in the *Heliconius* genus, with an independent loss in the *aoede* clade (four species that were previously classified as the *Neruda* genus) (Turner 1976; Beltrán et al. 2007; Kozak et al. 2015; Cicconardi et al., 2022). As *Heliconius* is the most speciose genus of the tribe, their novel ability to use pollen has likely contributed to their diversification, opening new niches to be exploited (*i. e*. through habitat partitioning, foodplant preference, foraging behaviour) and providing them with the energetic resources necessary for the maintenance of complex traits (Estrada and Jiggins 2002; Montgomery et al. 2016; Young & Montgomery, 2020; Couto et al. 2022). Indeed, the pollen-feeding behaviour of *Heliconius* butterflies has been associated with several aspects of their biology that diverge from the other heliconiine genera, including an elongated adult-lifespan (Dunlap-Pianka et al. 1977), prolonged fertility (Boggs et al. 1981; O’Brien et al. 2003), enlarged mushroom-bodies (Montgomery et al. 2016), foraging site fidelity (Moura, Corso, Montgomery, & Cardoso, 2022) and increased adult toxicity (de Castro et al. 2020)

Lepidopterans generally acquire most, if not all, of their nutrients during larval feeding. By supplying butterflies with amino acids, pollen feeding may have decoupled this partition (Boggs, 2009), providing a mechanism for further investment in adult behavioural strategies. Indeed, while most Lepidoptera tend to live relatively long lives as larvae and shorter lives as adults, *Heliconius* adults that have access to pollen can live up to 6 months, which is much longer than the regular average life-span of other heliconiines (∼1 month)(Brown, 1981), despite a similar larval period (Hebberecht, Melo-Flórez, Young, McMilllan, & Montgomery, 2022). Alongside this increased longevity, *Heliconius* butterflies also maintain their fecundity for longer than other heliconiines, such as *Dryas iulia*, showing limited evidence of reproductive senescence, unless deprived of pollen (Dunlap-Pianka et al., 1977). This prolonged fertility is energetically costly: a female butterfly can lay up to 9-18 eggs a day and they can live for many months, such that total resources allocated to oviposition exceed their own body mass. Indeed, O’Brien et al. (2003) used isotopic labelling to demonstrate the direct transfer of essential amino acid from pollen ingested by females to their eggs. Males also contribute to the cost of fertility by transferring nuptial gifts to the female during mating (Boggs and Gilbert, 1979; Cardoso, Roper, & Gilbert, 2009) which can exceed 5% of male body weight, and pollen resources may also be used for this purpose (Boggs, 1990). Although the relationship between diet, body weight, fertility and longevity seems obvious, there is a lack of empirical data about how pollen-feeding affects weight maintenance and how is this associated with the prolonged fertility of these butterflies.

Finally, the evolution of pollen feeding has also been associated with toxicity, a critical trait for chemically defended aposematic butterflies. *Heliconius* tend to have higher total concentrations of cyanogenic glucosides (CG) than other heliconiines (de Castro et al. 2019; Sculfort et al. 2020) and mature adults have higher concentrations than larvae and young adults (Nahrstedt and Davis 1983; de Castro et al. 2020). This is unusual in aposematic butterflies, which normally acquire their chemical defences from plants during larval feeding and therefore have more toxins as final instar larvae (Nishida, 2002). Whereas larvae of *Heliconius* balance between CG biosynthesis and sequestration from their obligatory Passifloraceae hostplants (de Castro et al. 2018; de Castro et al. 2021), adults can only biosynthesize these defence compounds, for which they need the amino acids valine and isoleucine (de Castro et al. 2020). It has been hypothesized that *Heliconius* butterflies use the essential amino acids from pollen for CG biosynthesis (Nahrstedt and Davis 1983). However, studies comparing the CG content of young *Heliconius* butterflies fed only sugar to those whose diet was supplemented with amino acids/pollen did not show any significant differences (Nahrstedt and Davis 1985; Cardoso and Gilbert 2013). This suggests that *Heliconius* butterflies might biosynthesize CGs initially using amino acids acquired during the larval stage, with resources from pollen-feeding only used later in adulthood.

Here, we explore how a dietary novelty can ease energetic constraints on life-history traits, using pollen feeding *Heliconius* as a case study. We investigate the effect of pollen-feeding on *H. erato* body weight, chemical defences, and fertility controlling for sex and age, and specifically comparing young adults (14d) with mature adults (45d). We therefore tested the hypothesis that mature butterflies that only had access to sugar during adulthood would have lower fertility, body weight and depleted chemical defences.

## Methods

### Rearing conditions of H. erato stock population

All experiments were performed using individuals from a stock population of *H. erato demophoon* kept at University of Cambridge. This population was sourced from Panamá city (Panama) and has been kept under insectary conditions for about 7 years. Adults were kept in breeding cages (60×60×90 cm) containing plants of *Passiflora biflora* for oviposition, as well as flowering *Lantana sp*. and few *Psiguria sp*. for adult feeding. Cages included feeders with artificial nectar made from 10% sucrose solution (m/v) with 1.5% (m/v) Vetark Critical Care Formula (CCF). *P. biflora* shoots with eggs were collected from the breeding cages and used to set up larval cages. Larvae were fed with fresh *P. biflora* shoots *ad libitium* until pupation. Larval cages were checked every other day and encountered pupae were transferred to pupal cages, where pupae were hung under a stick covered with a microfiber cloth Freshly emerged individuals in the pupal cages were transferred to breeding cages. All cages are kept at 25-28 °C, 60-80% humidity and 12h day/night cycle.

### Experimental Design and Diet treatments

Recently emerged adults (0-1 day after eclosion) were transferred to the experimental cages (60×60×90 cm). Only adults that had morphologically healthy, with uncrumpled dry wings were used in these experiments. In addition, only individuals with a forewings between 3.0 to 3.5 cm in length were used to control for potential size effects. One experimental cage was set up for each treatment (diet/age) and each had initially 8 males and 8 females (N=16). Butterflies that died during the first week of experiment were replaced to control for density. Butterflies were placed on feeders when added into the experimental cages to ensure that they would be able to find their food source. Each experimental cage received one of the diet treatments: 1) three feeders with artificial nectar made of 10% sucrose; or 2) three feeders with artificial nectar made of 10% sucrose + 1.5% amino acid supplement (CCF); or 3) three feeders with artificial nectar made of 10% sucrose and freshly collected *Lantana* flowers, as a natural source of pollen. Butterflies were fed *ad libitium*, with feeders and flowers were replaced every other day. Males and females in each treatment were allowed to mate freely. Experimental cages were kept for 14 days to assess the importance of amino acid on young butterflies and for 45 days to assess this effect on mature butterflies. All other heliconiines live for ∼1 month, therefore 45 days is the beginning of an adulthood period that is specific of mature *Heliconius* butterflies. All experimental cages were kept at the same environmental conditions used for husbandry (25-28 °C, 60-80% humidity and 12h day/night cycle). The protein concentration of *Lantana* pollen extracts and the CCF supplement was determined using the Pierce method (Supplementary Methods, Table S3).

### Fertility assays

At the end of the experiments, female butterflies were individually assayed for oviposition to evaluate the effect of the diet treatments on fertility, while males were kept in the experimental cages until sample collection. For the fertility assays, female butterflies were transferred into individual cages (30×30×40 cm) containing their previous diet (one feeder per cage, with one flower bouquet for the pollen treatment) and a *P. biflora* cutting with 5 expanded leaves for oviposition. After 48h of assay, eggs were counted, weighed, and collected for further analyses.

### Sample collection, metabolite extraction and HPLC-MS conditions

The weight of each butterfly was recorded at the end of the experiment (14 days or 45 days). 8 males and 8 females of freshly emerged butterflies (unfed, after 0-1 day of eclosion) were also weighed and collected as a baseline. Afterwards, butterflies were collected in 1 mL methanol 80% (v/v) for chemical analyses. All samples were kept at -20 °C until further processing. For the metabolite extraction, butterfly samples were homogenized (1mL methanol 80% (v/v)) using a porcelain mortar and pestle. Egg samples were homogenised in 300 μL methanol 80% (v/v) into their own collection tube using a small pestle. Extracts were centrifuged at 14,000 *g* for 5 min, filtered (45 μm) and collected for analyses in a LC-Orbitrap-MS/MS. LC-MS methods and analyses were conducted as described in de Castro et al. (2019). The *de novo* biosynthesized CGs linamarin, lotaustralin and epilotraustralin were quantified in the analysed samples, which had no other CGs. The absolute amount of each compound in each sample was calculated using the peak area of their sodium adduct applied to a regression curve stablished using pure standards. Raw chemical data as well as quantification methods can be found in https://doi.org/10.17863/CAM.92867)

### Statistical analyses

Statistical analyses and plots were performed in R. Shapiro-Wilko test was used to analyses if the variables were normally distributed (Table S1) and Levene’s test for the homogeneity of the variances (Table S2). ANOVA was used to evaluate the effect of diet, age and sex, as well as their interaction, on butterfly weight (Table 1, Table S3 for females only). Tukey HSD was used for pairwise comparisons between the different diet:age treatments in males and females. ANOVA was used to examine the effect of diet and age on CG per laid egg with Tukey HSD for pairwise comparisons. Kruskall-Wallis was used on variables that were not normally distributed: to analyse the effect of age and diet on laid eggs; and the effect of diet, age and sex on butterfly CG content.

**Table 1.** Effect of diet, age and sex on weight (grams per individual) of *H. erato* butterflies. The variables that have a significant effect on butterfly weight are marked in bold, with a * near their p value (p > 0.05). Diet treatments: sugar only, sugar + amino acid supplement, and sugar + pollen. Sex: female and male. Age: young (14d) and mature(45d).

| Variables | Three-way ANOVA |
| --- | --- |
| Diet | **$F_{2-95} = 4.597$, $p = 0.012^*$** |
| Age | **$F_{1-95} = 7.287$, $p = 0.008^*$** |
| Sex | **$F_{1-95} = 25.055$, $p = 2.5 \times 10^{-6}^*$** |
| Diet:Age | $F_{2-95} = 2.016$, $p = 0.139$ |
| Diet:Sex | $F_{2-95} = 0.486$, $p = 0.617$ |
| Age:Sex | **$F_{1-95} = 4.520$, $p = 0.036^*$** |
| Diet:Age:Sex: | $F_{2-95} = 1.791$, $p = 0.172$ |

## Results

### Females are more affected than males by the lack of pollen

Age, diet and sex significantly influenced the body weight of *H. erato* butterflies (Table 1). There was also a significant interaction between age and sex, which indicates that the body weight of males and females were differently affected through adulthood (Table 1). Indeed, overall females were heavier than males and they were more affected by the absence of amino acids on their adult diet (Figure 1). Mature females had lower weights by mid adulthood without access to nitrogen, but adult diet did not affect the weight of mature males (Figure 1). Mature females that had access to pollen were heavier than mature females fed sugar only, which had lower weights than freshly eclosed females (Figure 1). Males and females eclosed with similar weight (0d) (Figure S1).

**Figure 1.**
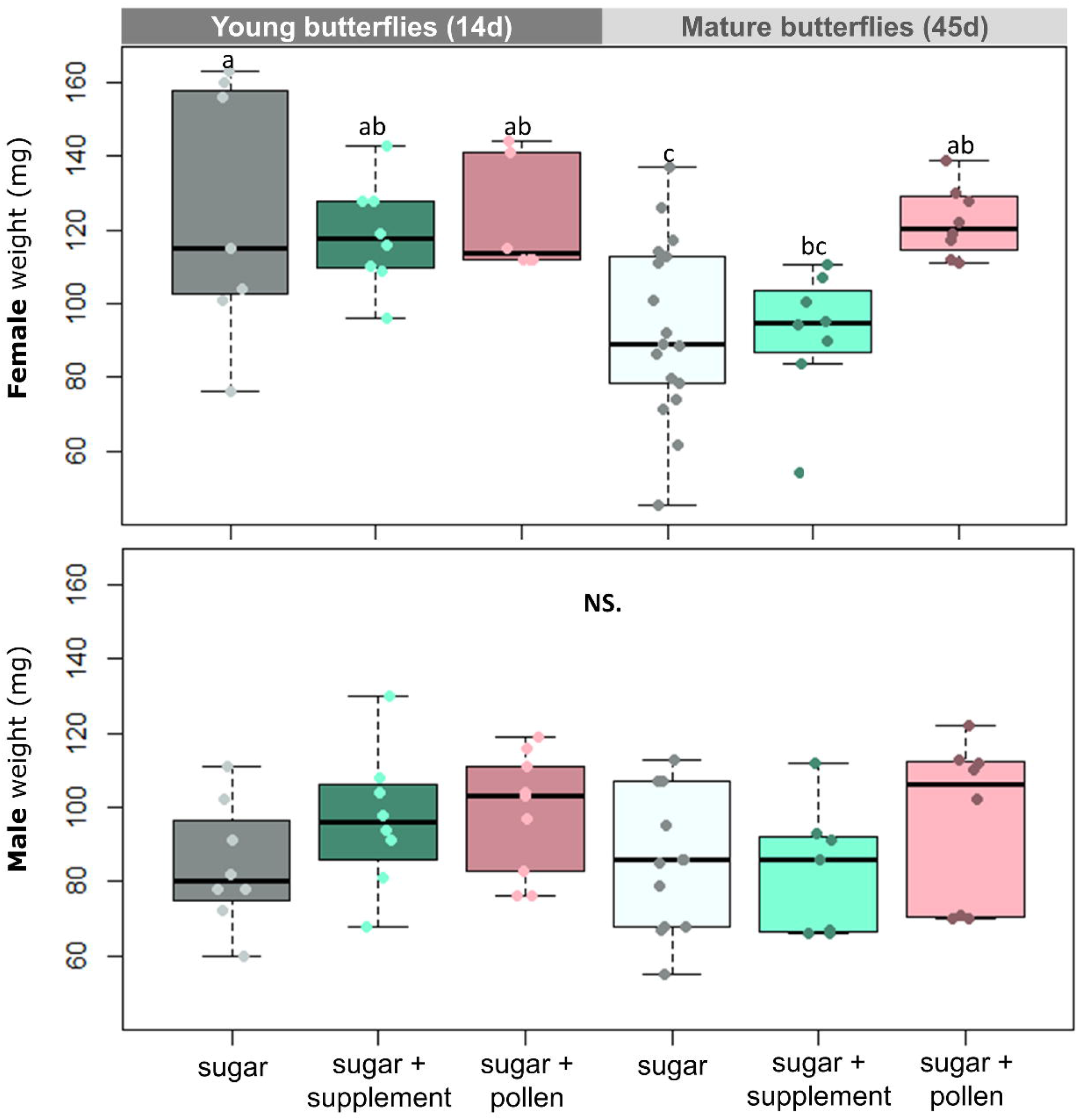
Effect of diet and age on the fresh weight of females (top) and males (bottom) of *H. erato*. Butterflies were fed either sugar or sugar + supplement (Critical Care Formula) or sugar + pollen (from Lantana flowers). Young butterflies were collected after 14d of trial while mature butterflies after 45d. Legend: Different letters over the boxplots correspond to statistically significant differences (Two-ways ANOVA, Tukey HSD). NS = not statistically significant (p> 0.05). Lines in the middle of boxplots correspond to the median and boxes to the lower and upper quartile. Dots correspond to values of each analysed replicate/individual butterfly.

### Access to pollen only affects the chemical defences of females

Males and females increased their CG content after eclosion (Figure S2) and kept their defences through adulthood, indicating that they intensively biosynthesize these compounds. Curiously, diet only affected the CG content of females (Figure 2. Kruskal-Wallis, Females: *X*^2^= 6.35, p= 0.048*; Males: *X*^2^= 2.115, p= 0.347), with butterflies having access to amino acids (supplement or pollen) showing greater CG content than those fed sugar alone. Young and mature butterflies of both sex had similar CG content (Figure 2. Kruskal-Wallis, Females: *X*^2^= 1.441, p= 0.23; Males: *X*^2^= 0.198, p= 0.656).

**Figure 2.**
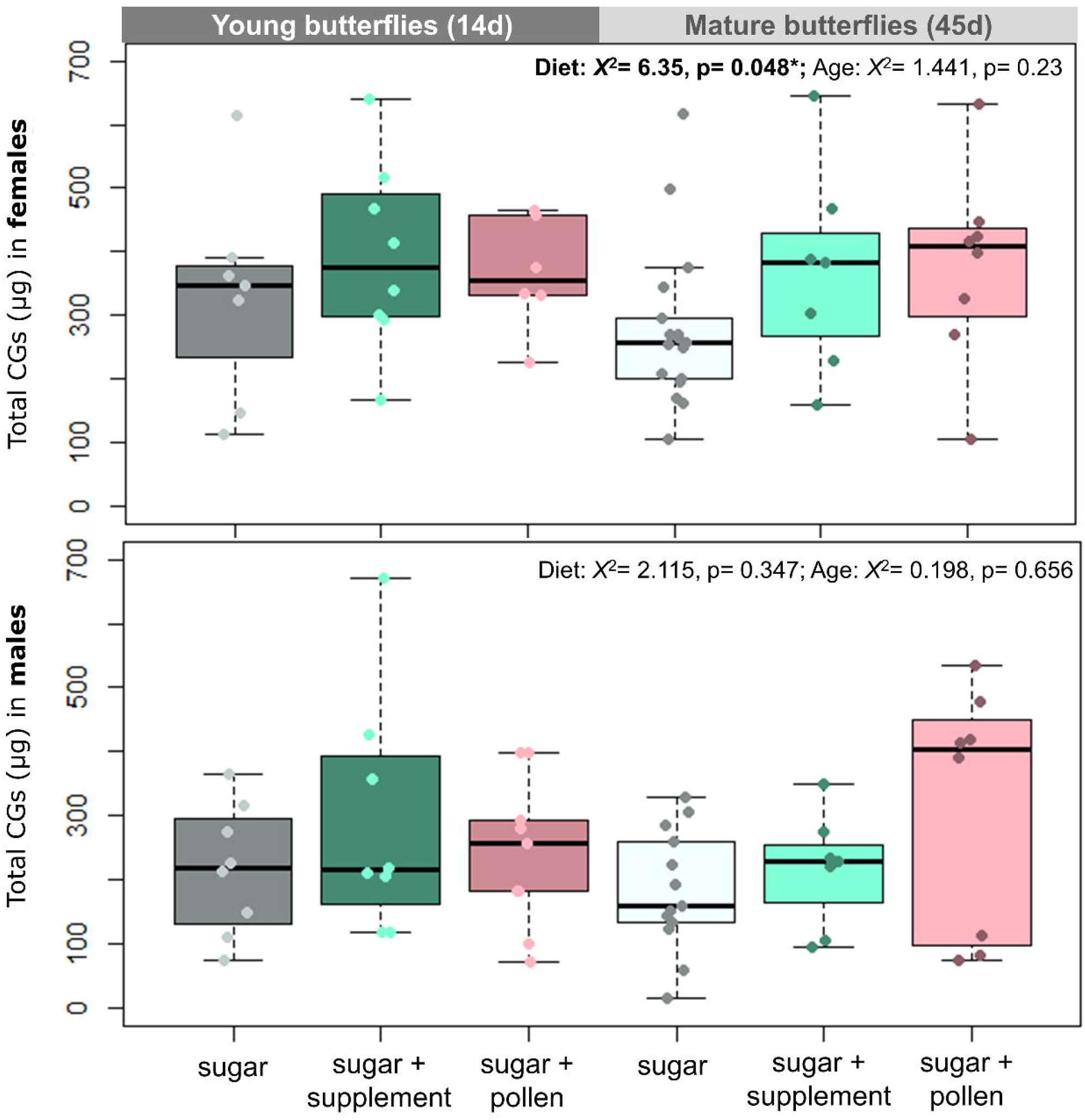
Effect of diet and age on cyanogenic glucosides content of females (top) and males (bottom) of *H. erato*. Butterflies were fed either sugar or sugar + supplement (Critical Care Formula) or sugar + pollen (from Lantana flowers). Young butterflies were collected after 14d of trial while mature butterflies after 45d. Statistical analyses on the top of the plots correspond to Kruskal-Willis on Diet and Age for the subsets. Lines in the middle of boxplots correspond to the median and boxes to the lower and upper quartile. Dots correspond to values of each analysed replicate/individual butterfly.

### Access to pollen delays reproductive senescence

Adult diet affected egg laying in mature butterflies of *H. erato* (Figure 3. Kruskal-Wallis, *X*^2^= 0.569, p= 0.017*), but not in young ones (Figure 3. Kruskal-Wallis, *X*^2^= 0.569, p= 0.752). Young females (14d) laid similar numbers of eggs regardless of their diet. In contrast, mature females (45d) that had access to pollen laid more eggs than butterflies that had access to sugar only, or sugar + supplement. This indicates that access to pollen delays reproductive senescence in *Heliconius*.

**Figure 3.**
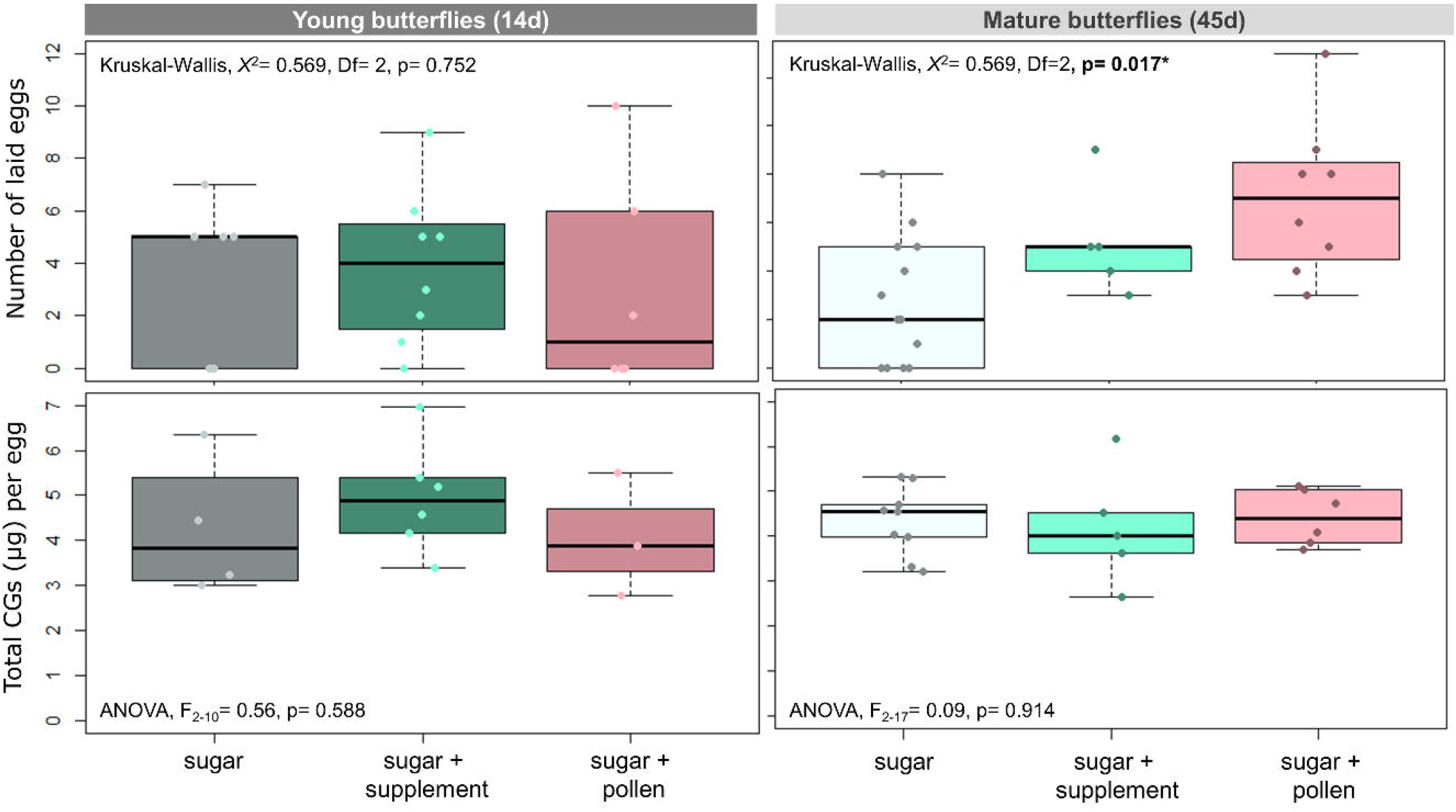
Number of laid eggs per female during fertility test (top) and cyanogenic glucose content per egg (bottom). Butterflies were fed either sugar or sugar + supplement (Critical Care Formula) or sugar + pollen (from *Lantana* flowers). Young butterflies were collected after 14d of trial while mature butterflies after 45d. Lines in the middle of boxplots correspond to the median and boxes to the lower and upper quartile. Dots correspond to values of each analysed replicate/total eggs laid by each butterfly.

In contrast to our expectations, adult diet did not affect parental allocation in the chemical defences (CG) of their eggs. Eggs of young (ANOVA, F_2-10_= 0.56, p= 0.588) and mature butterflies (ANOVA, F_2-17_= 0.09, p= 0.914) had similar concentrations of CG regardless of the diet of their parents.

### The nutritional uniqueness of pollen

Although the CCF supplement had far more proteins (651.70 ± 19.97 μg per mg DW) than the *Lantana* flower extract (pollen and nectar) (1.71 ± 0.45 μg per mg DW) (Table S3), it did not lead to improvements in the butterfly fitness traits. Thus, pollen might have an amino acid profile that fits better the nutritional needs of *Heliconius* and/or have them in a more acessible way (free amino acids instead of proteins/peptides).

## Discussion

### When does access to pollen start to be important and for whom?

Pollen-feeding eases energetic constraints and allows *Heliconius* to optimize multiple life-history traits simultaneously, aiding the maintenance of fertility, body weight and chemical defences during their prolonged adult-lifespan (Fig. 1, 2 and 3), as we hypothesized. Nevertheless, we found age-specific responses to this dietary novelty, as access to pollen has an effect on old butterflies (45d) of *H. erato*, but not on the young ones (14d). This is the first time to our knowledge that the effect of pollen deprivation on multiple life-history traits of *Heliconius* has been evaluated controlling for age. Studies supplementing the diets of other nectar-feeding long-lived nymphalids *(Polygonia c-album, Maniola jurtina*) with amino acids have not found an improvement in life-history traits (Karlsson and Wickman 1989; Grill et al. 2013). Our results therefore emphasize that adaptations were required to make use of pollen-derived amino acids in *Heliconius* butterflies (Dunlap-Pianka et al. 1977; Boggs et al. 1981).

The age-effects also reveal that the balance between larval and adult derived resources changes over the life course (Boggs, 2009). It is possible that the physiology of young *Heliconius* butterflies, including their fertility, initially relies mostly on resources acquired during larval feeding, as in the vast majority of butterflies and moths. Nevertheless, as *Heliconius* butterflies live relatively long adult lives, the reservoir of larval derived resources likely becomes depleted over time, such that the presence of amino acids in their adult diet becomes a crucial factor for the maintenance of the homeostasis. This is consistent with field data showing that older *Heliconius* butterflies generally collect more pollen then the young ones (Boggs et al., 1981; Boggs et al, 1990), which may indicate greater motivation as larval resources deplete.

This implies that studies evaluating the importance of pollen-feeding during adulthood for *Heliconius* butterflies likely need to be performed for periods longer than a month. Cardoso and Gilbert (2013) did not observed differences between the cyanide concentration from 20 day old *Heliconius* butterflies (*H. ethila, H. hecale* and *H. charithonia*) fed only sugar and supplemented with amino acids, as we observed here for females. The authors discussed the importance of larval diet shaping the chemical defences of young *Heliconius* butterflies. Additionally, the pheromone bouquet of 14 day old *Heliconius* males (genital and androconia) was also not affected by access to pollen during adulthood, only by the hostplant species used during larval feeding (Darragh et al., 2019). Indeed, the results seen here would probably be more striking if we have maintained the experiment for more than 45 days. Combined, these studies emphasize that the importance of the resources accumulated during larval feeding for young butterflies and the latter importance of pollen during adulthood.

It is a common knowledge among researchers breeding *Heliconius* under insectary conditions that they die sooner without access to pollen/amino acids in their adult diet. In this study, 45 days was not enough to observe differences in mortality between diet treatments (1-3 butterflies died in each treatment, mostly within the first week of eclosion), contrary to previous findings (Dunlap-Pianka et al. 1977). As previous experiments used *H. charithonia* (Dunlap-Pianka et al. 1977) while we used *H. erato*, this could indicate that different *Heliconius* species might rely on the amino acids acquired during adult-feeding sooner than others. Nevertheless, the previous work had few replicates (N=8 in total, N=3 after 40 days.) and might have underestimated how long *H. charithonia* can live on average without pollen. How different *Heliconius* species respond to the lack of pollen is an interesting question for the future.

### Do females pay a higher cost for reproduction than males when recourses are scarce?

This study demonstrates that access to amino acids delays reproductive senescence in *H. erato* females, as old females (45 d) supplemented with pollen lay as many eggs as young females (14d), whereas females fed only sugar lose fertility throughout adulthood (Fig. 3). Boggs (1990) observed that females of *H. charithonia* and *H. cydno* drastically increase pollen collection between 15-20 days old, possibly to maintain their fertility and chemical defences. This corroborates with the earlier work of Dunlap-Pianka et al. (1977) demonstrating that *H. charithonia* that have access to pollen can keep daily egg-laying rates until their natural death (up to 72d), however they continuously decrease their egg-production and reach ovarian depletion when pollen is absent.

The disparity between how females and males of the same species alter their life-history dynamics in response to resource availability has intrigued evolutionary ecologists. Without pollen-feeding, *H. erato* females lose weight (Fig. 1) and decrease their chemical defences (Fig. 2) as they get older, whereas males do not. As females collect significantly more pollen than males in the wild (Boggs et al., 1981), it could be that females feed more than males and therefore their fitness is more impacted by diet. Regardless, only females of *H. erato* were strongly affected by adult diet and this was reflected in their fertility, which might suggest that females are paying a higher energetic cost for reproduction than males when access to amino acids is limited. Even though diet did not affect male weight or CG content, we cannot discard the possibility that the effect of diet on fertility might be associated with other male fitness traits, such as sperm viability and quality of nuptial gifts (Boggs & Gilbert, 1979; Boggs, 1990).

Some of the old female butterflies in the cage supplemented with pollen had a strong smell of anti-aphrodisiac (personal observations) suggesting that they recently re-mated. Although re-mating was not expected in this experiment, since *H. erato* belongs to the monoandrous clade of *Heliconius* (Beltrán, Jiggins, Brower, Bermingham, & Mallet, 2007) and rarely re-mates in the wild (Cardoso et al. 2009; Walters et al. 2012), the insectary conditions might have induced them to re-mate. Re-mating would allow the transference of more nuptial gifts, which includes CGs, from the male to the female (Cardoso and Silva 2015), diluting the effect of pollen supplementation on male chemical defences and body weight. Further studies of spermatophore quality will be necessary to unravel the effect of pollen deprivation on the fitness of *Heliconius* males.

### Do high condition adults lay better protected eggs?

Many insects protect their eggs by transferring defensive compounds to them, which can improve offspring establishment. Thus, we hypothesized that butterflies with access to pollen would produce eggs with more CGs, as these compounds are not toxic when intact and can be stored in high concentrations. Old females of *H. erato* that had access to pollen are heavier (Fig. 1), had more CGs (Fig. 2) and laid more eggs (Fig. 3) than old females that had access to sugar only. Contrary to our predictions, this does not translate into a higher investment in the chemical defences of their offspring (Fig. 3). Eggs of butterflies from all ages and diets have similar CG content which suggests that this process is tightly regulated - butterflies might lay less or more eggs depending on their diet, but all eggs have a similar level of chemical defences. The amount of CG per egg observed here is similar to other heliconiines (Nahrstedt and Davis 1983; Nahrstedt and Davis 1985; Castro et al. 2020).

Our data demonstrate how strongly *H. erato* biosynthesize CG during adulthood to maintain their defences while also investing in the protection of their offspring, corroborating previous findings (Castro et al. 2020; Mattila et al. 2022). Considering that a *Heliconius* female lays ca. 10 eggs per day (Dunlap-Pianka et al. 1977), each egg has on average 3 μg of CG (Fig. 3) and they can live for 45 days. Egg-laying would therefore result in a depletion of over 1000 μg of CG from a female butterfly, which can be more than their whole reservoir of chemical defences at any one time (Fig. 3). In contrast, male contributuions for offspring chemical defences seems minimal (Cardoso & Gilbert, 2007). Pedigree experiments with *H. erato* also found strong maternal effects on offspring toxicity, but no paternal effects (Mattila et al., 2021).

Mattila et al. (2022) demonstrated that *Heliconius* butterflies keep their CG concentration at high levels during adulthood until their natural death. Indeed, if these aposematic butterflies lost their toxicity as they age, this would dilute the protection signal of their colour pattern. Thus, there is probably strong selection for *Heliconius* to maintain toxicity as they age, but it is likely challenging to maintain these levels while reproduction depletes their chemical reservoir (Fig. 2 and 3).

Moreover, valine and isoleucine are used as substrate for the biosynthesis of aliphatic CGs (Nahrstedt and Davis 1983). These are essential amino acids that have to be acquired by diet (not produced by animals) (O’Brien et al., 2002) and they tend to be abundant in pollen (Gilbert 1972). This suggests a strong effect of pollen-feeding on chemical defences in *Heliconius*. Yet, a lack of pollen/amino acids during adult-feeding does not affect the chemical defences of young *Heliconius* butterflies. As already discussed, access to pollen would become crucial at later stages of adulthood, but the remaining question is: where did the valine and isoleucine used for CG biosynthesis come from during the first weeks of *Heliconius* adulthood in the control group (sugar only)? A recent comparative genomic study has found that two hexamerins, storage proteins, have been duplicated multiple times in heliconiines (Cicconardi et al., 2022). Hexamerins might provide valine and isoleucine for CG biosynthesis during the beginning of their adulthood, if pollen is not available. Moreover, valine and isoleucine might be produced by bacteria in the microbiome of these butterflies, as happens for other insects (Jing, Qi, & Wang, 2020), a hypothesis that can be investigated in the future.

In summary, although the link between pollen-feeding, fertility and chemical defences in *Heliconius* butterflies is clear, these interactions are more complex than initially predicted. We demonstrated that there is an age-specific and sex-specific effect of pollen-feeding on life-history traits. Older females supplemented with pollen were heavier, more toxic and laid more eggs than those in the control diet, suggesting that this dietary innovation has eased energetic constraints and led to optimization of multiple life-history traits.

## Supporting information

Dataset on diet protein content

Metadata

## DATA ACCESSIBILITY

The metadata and coding script associated with this publication are available as supplementary material. These files and the raw chemical data are available in the Apollo repository from University of Cambridge at https://doi.org/10.17863/CAM.92867.

## REFERENCES

Beltrán, M., Jiggins, C. D., Brower, A. V. Z., Bermingham, E., & Mallet, J. (2007). Do pollen feeding, pupal-mating and larval gregariousness have a single origin in Heliconius butterflies? Inferences from multilocus DNA sequence data. Biological Journal of the Linnean Society, 92(2), 221–239. https://doi.org/10.1111/J.1095-8312.2007.00830.X

Boggs, C. L. & Gilbert, L. E. (1979). Male contribution to egg production in butterflies: evidence for transfer of nutrients at mating. Science 206(4414), 84–84. https://doi.org/10.1126/science.206.4414.83.

Boggs, C. L., Smiley, J. T., & Gilbert, L. E. (1981). Patterns of pollen exploitation by Heliconius butterflies. Oecologia 48:2, 48(2), 284–289. https://doi.org/10.1007/BF00347978

Boggs, C. L. (1981b). Nutritional and Life-History Determinants of Resource Allocation in Holometabolous Insects. The American Naturalist, 117(5), 692–709. https://doi.org/10.1086/283753

Boggs, C. L. (2009). Understanding insect life histories and senescence through a resource allocation lens. Functional Ecology, 23(1), 27–37. https://doi.org/10.1111/J.1365-2435.2009.01527.X

Boggs, C., & Iyengar, V. (2022). Age-specific and Sex-specific nectar and pollen use by a butterfly pollinator. bioRxiv, 2022-05. https://doi.org/10.1101/2022.05.19.492749

Brown, K. S. (1981). The Biology of Heliconius and Related Genera. Ann. Rev. EntomoL, 26(155), 427–6. Retrieved from http://www.annualreviews.org/aronline

Cardoso, M. Z., & Gilbert, L. E. (2013). Pollen feeding, resource allocation and the evolution of chemical defence in passion vine butterflies. Journal of Evolutionary Biology, 26(6), 1254–1260. https://doi.org/10.1111/jeb.12119

Cardoso, Marcio Z., & Silva, E. S. (2015). Spermatophore Quality and Production in two Heliconius Butterflies with Contrasting Mating Systems. Journal of Insect Behavior, 28(6), 693–703. https://doi.org/10.1007/s10905-015-9536-y

Cardoso, Márcio Zikán, & Gilbert, L. E. (2007). A male gift to its partner? Cyanogenic glycosides in the spermatophore of longwing butterflies (Heliconius). Die Naturwissenschaften, 94(1), 39–42. https://doi.org/10.1007/s00114-006-0154-6

Cardoso, Márcio Zikán Roper, J. J., & Gilbert, L. E. (2009). Prenuptial agreements: mating frequency predicts gift-giving in Heliconius species. Entomologia Experimentalis et Applicata, 131(2), 109–114. https://doi.org/10.1111/J.1570-7458.2009.00837.X

Cicconardi, F., Milanetti, E., Pinheiro De Castro, É. C., Mazo-Vargas, A., Van Belleghem, S. M., Ruggieri, A. A., … Piedras, R. (2022). Evolutionary dynamics of genome size and content during the adaptive radiation of Heliconiini butterflies. BioRxiv. https://doi.org/10.1101/2022.08.12.503723

Couto, A., Young, F. J., Atzeni, D., Marty, S., Melo-Florez, L., Hebberecht, L., … & Montgomery, S. H. (2022). Rapid expansion and visual specialization of learning and memory centers in Heliconiini butterflies. bioRxiv, 2022-09. https://doi.org/10.1101/2022.09.23.509163

Darragh, K., Byers, K. J. R. P., Merrill, R. M., McMillan, W. O., Schulz, S., & Jiggins, C. D. (2019). Male pheromone composition depends on larval but not adult diet in Heliconius melpomene. Ecological Entomology, 44(3), 397. https://doi.org/10.1111/EEN.12716

de Castro Érika C.P., Musgrove, J., Bak, S., McMillan, W. O., & Jiggins, C. D. (2021). Phenotypic plasticity in chemical defence of butterflies allows usage of diverse host plants. Biology Letters, 17(3), 20200863. https://doi.org/10.1098/rsbl.2020.0863

de Castro Érika C. P., Zagrobelny, M., Zurano, J. P., Cardoso, M. Z., Feyereisen, R., & Bak, S. (2019). Sequestration and biosynthesis of cyanogenic glucosides in passion vine butterflies and consequences for the diversification of their host plants. Ecology and Evolution, 9(9), 5079–5093. https://doi.org/10.1002/ece3.5062

de Castro, É. C. P., Zagrobelny, M., Cardoso, M. Z., & Bak, S. (2018). The arms race between heliconiine butterflies and Passiflora plants – new insights on an ancient subject. Biological Reviews, 93(1), 555–573. https://doi.org/10.1111/brv.12357

de Castro, É. C. P., Demirtas, R., Orteu, A., Olsen, C. E., Motawie, M. S., Zikan Cardoso, M., Zagrobelny, M., & Bak, S. (2020). The dynamics of cyanide defences in the life cycle of an aposematic butterfly: Biosynthesis versus sequestration. Insect Biochemistry and Molecular Biology, 116, 103259. https://doi.org/10.1016/j.ibmb.2019.103259

Dunlap-Pianka, H., Boggs, C. L., & Gilbert, L. E. (1977). Ovarian dynamics in heliconiine butterflies: Programmed senescence versus eternal youth. Science, 197(4302), 487–490. https://doi.org/10.1126/SCIENCE.197.4302.487

Estrada, C., & Jiggins, C. D. (2002). Patterns of pollen feeding and habitat preference among Heliconius species. Ecological Entomology, 27(4), 448–456. https://doi.org/10.1046/j.1365-2311.2002.00434.x

Gilbert, L. E. (1972). Pollen feeding and reproductive biology of Heliconius butterflies. Proceedings of the National Academy of Sciences of the United States of America, 69(6), 1403–1407. https://doi.org/10.1073/PNAS.69.6.1403

Grill, A., Cerny, A., & Fiedler, K. (2013). Hot summers, long life: egg laying strategies of Maniola butterflies are affected by geographic provenance rather than adult diet. Contributions to Zoology, 82(1), 27–36. https://doi.org/10.1163/18759866-08201002

Harpel, D., Cullen, D. A., Ott, S. R., Jiggins, C. D., & Walters, J. R. (2015). Pollen feeding proteomics: Salivary proteins of the passion flower butterfly, Heliconius melpomene. Insect Biochemistry and Molecular Biology, 63, 7–13. https://doi.org/10.1016/J.IBMB.2015.04.004

Healy, K., Ezard, T. H. G., Jones, O. R., Salguero-Gómez, R., & Buckley, Y. M. (2019). Animal life history is shaped by the pace of life and the distribution of age-specific mortality and reproduction. Nature Ecology & Evolution 2019 3:8, 3(8), 1217–1224. https://doi.org/10.1038/s41559-019-0938-7

Hebberecht, L., Melo-Flórez, L., Young, F. J., McMillan, W. O., & Montgomery, S. H. (2022). The evolution of adult pollen feeding did not alter postembryonic growth in Heliconius butterflies. Ecology and Evolution, 12(6). https://doi.org/10.1002/ECE3.8999

Hikl, A. L., & Krenn, H. W. (2011). Pollen processing behavior of heliconius butterflies: A derived grooming behavior. Journal of Insect Science, 11(1). https://doi.org/10.1673/031.011.9901

Jing, T. Z., Qi, F. H., & Wang, Z. Y. (2020). Most dominant roles of insect gut bacteria: Digestion, detoxification, or essential nutrient provision? Microbiome, 8(1). https://doi.org/10.1186/s40168-020-00823-y

Johnson, S. A., & Nicolson, S. W. (2001). Pollen digestion by flower-feeding scarabaeidae: Protea beetles (Cetoniini) and monkey beetles (Hopliini). Journal of Insect Physiology, 47(7), 725–733. https://doi.org/10.1016/S0022-1910(00)00166-9

Karlsson, B., & Wickman, P.-O. (1989). The Cost of Prolonged Life: An Experiment on a Nymphalid Butterfly. Source: Functional Ecology, 3(4), 399–405.

Kozak, K. M., Wahlberg, N., Neild, A. F. E., Dasmahapatra, K. K., Mallet, J., & Jiggins, C. D. (2015). Multilocus species trees show the recent adaptive radiation of the mimetic Heliconius butterflies. Systematic Biology, 64(3), 505–524. https://doi.org/10.1093/sysbio/syv007

Krenn, H. W. (2008). Feeding behaviours of neotropical butterflies (Lepidoptera, Papilionoidea). Biologiezentrum Linz/Austria, 80, 295–304.

Krenn, H. W., Eberhard, M. J. B., Eberhard, S. H., Hikl, A.-L., Huber, W., & Gilbert, L. E. (2009). Mechanical damage to pollen aids nutrient acquisition in Heliconius butterflies (Nymphalidae). Arthropod-Plant Interactions, 3(4), 203–208. https://doi.org/10.1007/s11829-009-9074-7

Law, R. (1979). Optimal Life Histories Under Age-Specific Predation. The American Naturalist, 114(3), 399–417. https://doi.org/10.1086/283488

Mattila, A. L. K., Jiggins, C. D., Opedal, Ø. H., Montejo-Kovacevich, G., De Castro, É. C. P., McMillan, W. O., … Saastamoinen, M. (2021). Evolutionary and ecological processes influencing chemical defense variation in an aposematic and mimetic Heliconius butterfly. PeerJ, 9, e11523. https://doi.org/10.7717/peerj.11523

Mattila, A. L. K., Jiggins, C. D., & Saastamoinen, M. (2022). Condition dependence in biosynthesized chemical defenses of an aposematic and mimetic Heliconius butterfly. Ecology and Evolution, 12(6). https://doi.org/10.1002/ece3.9041

Montgomery, S. H., Merrill, R. M., & Ott, S. R. (2016). Brain composition in Heliconius butterflies, posteclosion growth and experience-dependent neuropil plasticity. Journal of Comparative Neurology, 524(9), 1747–1769. https://doi.org/10.1002/cne.23993

Moura, P. A., Corso, G., Montgomery, S. H., & Cardoso, M. Z. (2022). True site fidelity in pollenfeeding butterflies. Functional Ecology, 36(3), 572–582. https://doi.org/10.1111/1365-2435.13976

Nahrstedt, a., & Davis, R. H. (1985). Biosynthesis and quantitative relationships of the cyanogenic glucosides, linamarin and lotaustralin, in genera of the Heliconiini (Insecta: Lepidoptera). Comparative Biochemistry and Physiology Part B: Comparative Biochemistry, 82(4), 745–749. https://doi.org/10.1016/0305-0491(85)90519-X

Nahrstedt, A., & Davis, R. H. (1983). Occurrence, variation and biosynthesis of the cyanogenic glucosides linamarin and lotaustralin in species of the Heliconiini (Insecta: Lepidoptera). Comparative Biochemistry and Physiology -- Part B: Biochemistry And, 75(1), 65–73. https://doi.org/10.1016/0305-0491(83)90041-X

Nishida, R. (2002). Sequestration of defensive substances from plants by Lepidoptera. Annual Review of Entomology, 57–92.

O’Brien, D. M., Fogel, M. L., & Boggs, C. L. (2002). Renewable and nonrenewable resources: Amino acid turnover and allocation to reproduction in Lepidoptera. Proceedings of the National Academy of Sciences, 99(7), 4413–4418. https://doi.org/10.1073/pnas.072346699

O’Brien, D. M., Boggs, C. L., & Fogel, M. L. (2003). Pollen feeding in the butterfly Heliconius charitonia: Isotopic evidence for essential amino acid transfer from pollen to eggs. Proceedings of the Royal Society B: Biological Sciences, 270(1533), 2631–2636. https://doi.org/10.1098/rspb.2003.2552

Sculfort, O., Castro, E. C. P., Kozak, K. M., Bak, S., Elias, M., Nay, B., & Llaurens, V. (2020). Variation of chemical compounds in wild Heliconiini reveals ecological factors involved in the evolution of chemical defenses in mimetic butterflies. Ecology and Evolution, 10(5), 2677–2694. https://doi.org/10.1002/ece3.6044

Smith, G., Kelly, J. E., Macias-Muñoz, A., Butts, C. T., Martin, R. W., & Briscoe, A. D. (2018). Evolutionary and structural analyses uncover a role for solvent interactions in the diversification of cocoonases in butterflies. Proceedings of the Royal Society B: Biological Sciences, 285(1870). https://doi.org/10.1098/RSPB.2017.2037

Smith, G., Macias-Muñoz, A., & Briscoe, A. D. (2016). Gene Duplication and Gene Expression Changes Play a Role in the Evolution of Candidate Pollen Feeding Genes in Heliconius Butterflies. Genome Biology and Evolution, 8(8), 2581–2596. https://doi.org/10.1093/gbe/evw180

Swanson, E. M., Espeset, A., Mikati, I., Bolduc, I., Kulhanek, R., White, W. A., … Snell-Rood, E. C. (2016). Nutrition shapes life-history evolution across species. Proceedings of the Royal Society B: Biological Sciences, 283(1834). https://doi.org/10.1098/rspb.2015.2764

Turner, J. R. G. (1976). Adaptive radiation and convergence in subdivisions of the butterfly genus Heliconius (Lepidoptera: Nymphalidae). Zoological Journal of the Linnean Society, 58(4), 297–308. https://doi.org/10.1111/j.1096-3642.1976.tb01000.x

Wäckers, F. L., Romeis, J., & Van Rijn, P. (2007). Nectar and pollen feeding by insect herbivores and implications for multitrophic interactions. Annual Review of Entomology, 52, 301–323. https://doi.org/10.1146/annurev.ento.52.110405.091352

Walters, J. R., Corbins, C., Hardcastle, T. J., & Jiggins, C. D. (2012). Evaluating female remating rates in light of spermatophore degradation in Heliconius butterflies: pupal-mating monandry versus adult-mating polyandry. Ecological Entomology, 37(4), 257–268. https://doi.org/10.1111/J.1365-2311.2012.01360.X

Young, F. J., & Montgomery, S. H. (2020). Pollen feeding in Heliconius butterflies: the singular evolution of an adaptive suite. Proceedings of the Royal Society B, 287(1938). https://doi.org/10.1098/RSPB.2020.1304

