## Supplementary Figure and Tables for "Pollen-feeding delays reproductive senescence and maintains toxicity of *Heliconius erato*"

### SUPPLEMENTARY MATERIAL

#### Supplementary Methods

##### *Sample preparation and protein quantification*

Pollen and nectar samples of *Lantana sp* and *Psiguria sp* were collected by pipetting and resuspending ultra-pure water into their flowers (10 µL per flower) multiple times. Three pooled replicates were collected per species, each composed of the exudate of 10 flowers for *Psiguria* and 20 flowers for *Lantana*. For the supplement, 20 mg Critical Care Formula (CCF) was mixed in 10 mL ultra-pure water per replicate. The protein assay was performed using the Pierce$^{TM}$660 nm method (ThermoFisher) following the manufacturer's instruction: 100 µL of each sample was incubated into 1.5 mL of Pierce Reagent and their absorbances read at 660 nm against a blank (100 µL ultra-pure water and 1.5 mL Pierce Reagent). A standard curve composed of Bovine Serum Albumin solutions at different concentrations was also assayed and the established regression equation used to calculate the amount of protein per sample. The remaining flower samples were dried in a SpeedVac (medium speeds) for 2 hours to establish their dry mass.

#### Supplementary Tables

**Table S1.** Normality test (Shapiro-Wilk) on the analysed variables. * corresponds to variables that have a normal distribution (p > 0.05).

| Variables | Shapiro-Wilk |
|---|---|
| Butterfly weight (mg) | **W= 0.987, p= 0.306*** |
| Laid eggs | W= 0.926, p= 0.0056 |
| Butterfly CG content (ng per butterfly) | W= 0.946, p= 0.0001 |
| Egg CG content (ng per egg) | **W= 0.972, p= 0.668*** |

**Table S2.** Homogeneity of the variances (Levene's Test) on the analysed variables, controlling by diet, age and sex. * corresponds to variables where the homogeneity assumption is met (p > 0.05).

| Variables | Levene's Test |
|---|---|
| Butterfly weight (mg) | **$F_{11\text{-}95}$= 1.468, p= 0.157*** |
| Laid eggs | **$F_{5\text{-}41}$= 0.363, p= 0.871*** |
| Butterfly CG content (ng per butterfly) | **$F_{11\text{-}94}$= 0.628, p= 0.801*** |
| Egg CG content (ng per egg) | **$F_{5\text{-}27}$= 0.636 p= 0.674*** |

**Table S3.** Protein concentration (µg/mg DW) in supplement and flowers extracts used as an amino acid source to feed *Heliconius* butterflies.

| Amino Acid Source | Protein Concentration (µg/mg DW) |
|---|---|
| CCF supplement | 651.70 ± 19.97 |
| *Lantana sp* pollen and nectar extract | 1.71 ± 0.45 |
| *Psiguria sp* pollen and nectar extract | 9.20 ± 0.97 |

#### Supplementary Figures

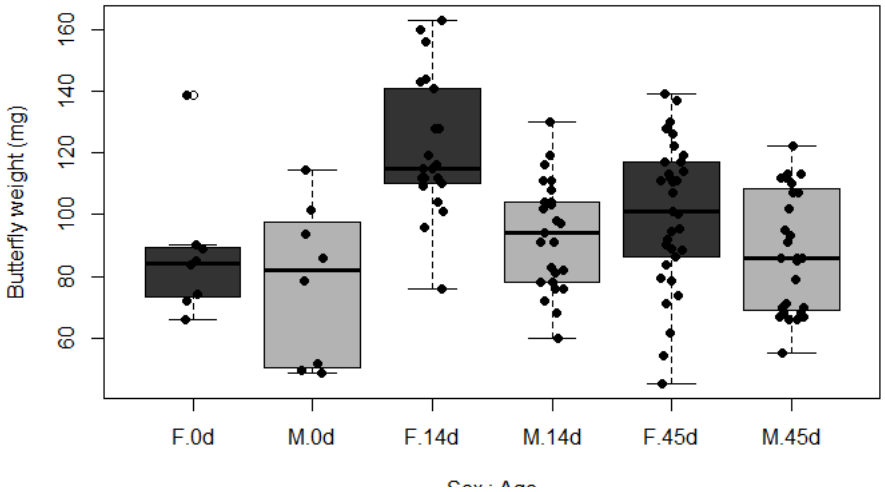


**Figure S1.** Weight of *H. erato* females (F) and males (M) at different ages (0d, 14d and 45d after eclosion).

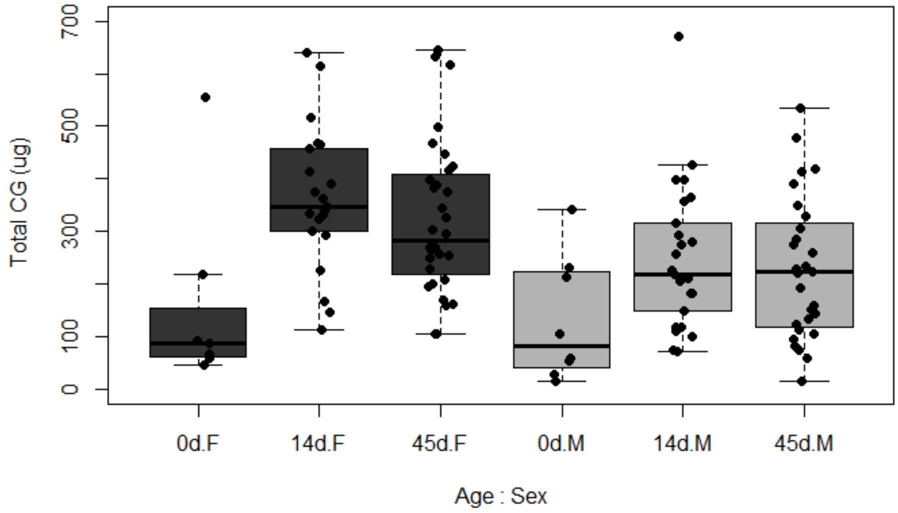


**Figure S2.** Total content of cyanogenic glucosides (CG) in females (F) and males (M) of *H. erato* at different ages (0d, 14d, and 45d after eclosion)

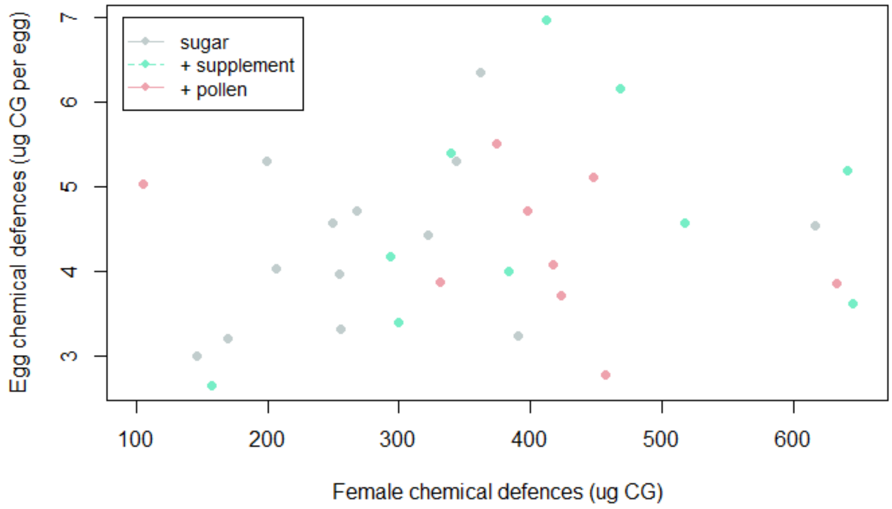


**Figure S3.** Relationship between egg chemical defences and female chemical defences. The correlation is not significant ($R^2$= 0.033, $F_{1\text{-}30}$= 1.021, p= 0.32).
